# SARS-CoV-2 infection induces dopaminergic neuronal loss in midbrain organoids during short and prolonged cultures

**DOI:** 10.1101/2023.03.20.533485

**Authors:** Javier Jarazo, Eveline Santos da Silva, Enrico Glaab, Danielle Perez-Bercoff, Jens C. Schwamborn

**Affiliations:** Developmental and Cellular Biology, Luxembourg Centre for Systems Biomedicine University of Luxembourg, 7 avenue des Hauts-Fourneaux, L-4362, Esch-sur-Alzette, Luxembourg; OrganoTherapeutics SARL, 6A, avenue des Hauts-Fourneaux, L-4365 Esch-sur-Alzette, Luxembourg; HIV Clinical and Translational Research Group, Department of Infection and Immunity, Luxembourg Institute of Health, 29 rue Henri Koch, L-4354, Esch-sur-Alzette, Luxembourg; Biomedical Data Science, Luxembourg Centre for Systems Biomedicine University of Luxembourg, 7 avenue des Hauts-Fourneaux, L-4362, Esch-sur-Alzette, Luxembourg

## Abstract

COVID-19 is mainly associated with respiratory symptoms, although several reports showed that SARS-CoV-2 affects the nervous system. We evaluated the effects of infection in prolonged culture of midbrain organoids, showing that the virus induces changes in gene expression, and fragmentation and loss of dopaminergic neurons. Our findings highlight the direct viral-induced damage to midbrain organoids indicating the relevance of assessing the neurological long-term evolution of COVID-19 patients.

## Main text

The ongoing COVID-19 pandemic caused by SARS-CoV-2 can cause acute symptoms ranging from a simple cold to respiratory insufficiency and cytokine storm that can lead to death^1^. Patients can also develop chronic symptoms including respiratory and cognitive impairment, such as fatigue, depression, and inability to focus, a condition termed Post-Acute Sequelae of SARS-CoV-2 infection (PASC), or “long Covid”^2^. Recent evidence showed that SARS-CoV-2 not only affects the respiratory tract, but other systems as well, such as the central nervous system, albeit with no major cytopathological alterations in the brain after autopsies^3^. Other post-mortem reports indicated that although generally mild signs were detected in the overall brain, the area that showed a clear inflammatory process was the brainstem^4, 5^. This region, consisting of Medulla, Pons and Midbrain, was also identified in a longitudinal study in several COVID-19 patients (mostly with mild symptoms or asymptomatic) presenting a reduction in volume detected by magnetic resonance imaging (MRI)^6^. Considering these observations together with the nuclei localization of major neuronal systems regulating critical body functions^7^, we planned to evaluate in vitro, long-term infection-related changes in the midbrain, and infer parallels with a neurodegenerative condition like Parkinson’s disease (PD).

Modelling COVID-19 in vitro using organoids has shed some light on the changes induced by SARS-CoV-2 infection^8^. Several studies based on brain organoids mimicking the overall central nervous system (CNS) highlight the neural tropism of the virus and describe a range of impacts extending from reduced inflammatory responses to cell death^9^. These studies focused on the early responses to viral infection (longest period evaluated was 72h post-infection^9, 10^) and used organoids that do not have a comparable proportion of dopaminergic neurons as the one present in the midbrain. In contrast, the model used in this study recapitulates the main characteristics of the midbrain with functional activity such as synapse formation, spontaneous firing, and dopamine release^11^. It also shows hallmarks of PD like reduction of dopaminergic neurons, when derived from individuals presenting a genetic form of the disease^12, 13^. The aim of this study was to determine if SARS-CoV-2 has the ability to infect midbrain organoids, to assess the direct effect of the virus on dopaminergic neurons and astrocytes, and to detect if the virus can induce a neurodegenerative process after an extended culture time.

Midbrain organoids were exposed to 0.05 moi of SARS-CoV-2 for 16 hours and changes post-infection were analyzed after short- and long-term cultures of 4- and 28-days post-infection (dpi) respectively. Using an automated image analysis platform (Extended Data Fig. 1a), features of the cell types present in the midbrain organoids were extracted. At 4 dpi, a positive signal for the SARS-CoV-2 nucleocapsid (N) was detected not only at the external boundaries but also in the inner parts of the organoid, while no signal was seen in the mock condition (Fig. 1a). A different antibody confirmed this observation (Extended Data Fig. 1b). Even though the level of colocalization between dopaminergic neurons (identified by tyrosine hydroxylase, TH) and SARS-CoV-2 was high, not all TH+ neurons stained positive for N (Fig. 1a). Infection with SARS-CoV-2 did not increase the proportion of pyknotic nuclei when normalized to the total amount of nuclei signal (Fig. 1b,c). There was a decrease in N-staining pixels over time, suggesting non-productive infection or reflecting cell-death (Fig. 1c). The levels of dopaminergic neurons were significantly reduced at 4 and 28 dpi. Although the amount of TH+ pixels recovered over time, it was not to the levels observed in the untreated condition (Fig. 1c). When assessing the degree of fragmentation of TH+ neurons, an early sign of degeneration, we observed that over time TH+ neurons presented a significantly increased disruption of neurite continuity (Fig. 1b,c). Within the same infected organoid, TH+ and SARS-CoV-2 positive neurons presented an altered morphology and high degree of neurite fragmentation compared to uninfected TH+ neurons (Fig. 1b). Altogether, these observations suggest that SARS-CoV-2 infects TH+ cells and leads to cell death.

**Fig. 1.**
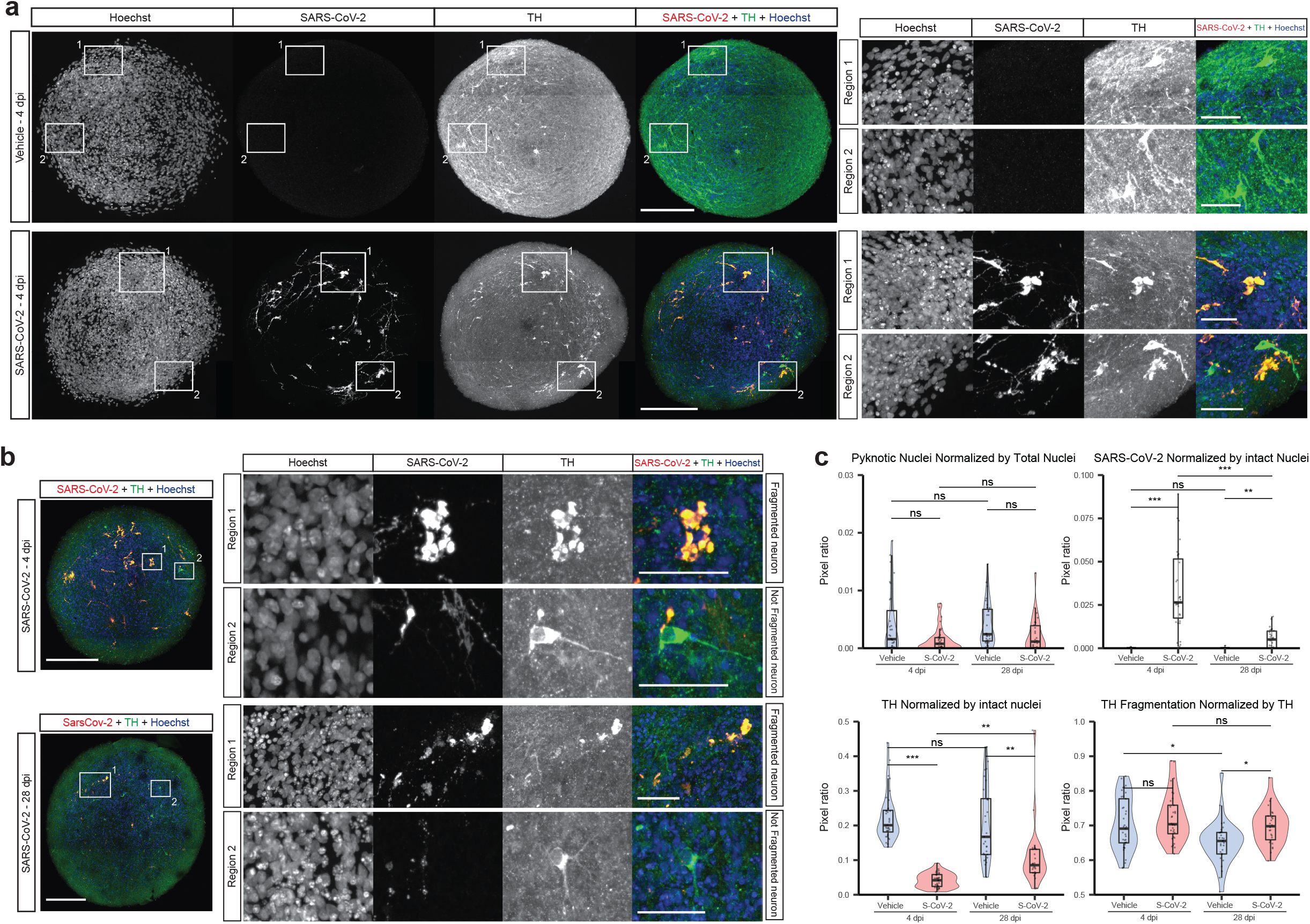
SARS-CoV-2 colocalizes with TH inducing neuronal fragmentation. **a**, Representative images of midbrain organoid sections stained for SARS-CoV-2 Nucleocapsid (N), Tyrosine Hydroxylase (TH) and a nuclear dye (Hoechst) for the short-term culture (4 dpi) and mock condition. Separate channel images are presented in grey scale, combined images with SARS-CoV-2 N in red, TH in green and Hoechst in blue. Numbered boxes represent zoomed regions in the panel to the right. Scale bar in main panel, 200μm. Scale bar in zoomed region, 50 μm. **b**, Representative images of midbrain organoid sections stained for SARS-CoV-2 N, TH and Hoechst for the short- and long-term cultures (4 dpi and 28 dpi respectively). Separate channel images are presented in grey scale, combined images with SARS-CoV-2 N in red, TH in green and Hoechst in blue. Numbered boxes represent zoomed regions of fragmented and unfragmented dopaminergic neurons. Scale bar in main panel, 200μm. Scale bar in zoomed region, 50 μm. **c**, Quantification of pixels of the respective masks normalized to a specific marker in the different conditions, Vehicle (V) or SARS-CoV-2 infection (S), at different timepoints, 4 dpi or 28 dpi; *n*(V4/S4/V28/S28): 37/33/31/26 sections, from 4 organoids per group. Top-left panel, represents the pixel ratio between pyknotic nuclei and total nuclei. Kruskal-Wallis (*P*<0.01) followed by Dunn’s test adjusted by Benjamini-Hochberg. Top-right panel, represents the pixel ratio between SARS-CoV-2 N and intact (not pyknotic) nuclei. Kruskal-Wallis (*P*<0.0001) followed by Dunn’s test adjusted by Benjamini-Hochberg. Bottom-left panel, represents the pixel ratio between TH and intact nuclei. Kruskal-Wallis (*P*<0.0001) followed by Dunn’s test adjusted by Benjamini-Hochberg. Bottom-right panel, represents the pixel ratio between fragmented TH signal and total TH signal. Kruskal-Wallis (*P*<0.001) followed by Dunn’s test adjusted by Benjamini-Hochberg. Individual data points are shown with their distribution and boxplots representing the 25^th^ (lower hinge), median (thick line), and 75^th^ quartiles. Whiskers represent 1.5*IQR (inter-quartile range). \**P*<0.05, \*\**P*<0.01, \*\*\**P*<0.001, and ns (not significant) *P*>0.05.

We then evaluated whether SARS-CoV-2 can infect astrocytes, by assessing the expression of GFAP and S100b^13, 14^. We observed that while their levels and colocalization were reduced during the early stages of infection, they increased over time, with S100b remaining significantly lower than control conditions (Fig. 2a,b). SARS-CoV-2 colocalized significantly more with certain types of cells (Fig. 2c-e). When assessing the type of cell that presented the highest proportion of N signal, neurons positive for the protein MAP2 were the most affected in both short- and long-term cultures (Fig. 2d). However, when looking at the proportion of a particular cell type that stained positive for SARS-CoV-2, dopaminergic neurons were the most affected, reaching around 40% of TH+ neurons in the midbrain organoid showing positive signal for the virus in short-term cultures (Fig. 2d). A significantly higher infection of GFAP+ and TH+ cells (when normalized to their respective abundance and the level of SARS-CoV-2 signal in the organoid) was observed for both short- and long-term cultures (Fig. 2e).

**Fig. 2.**
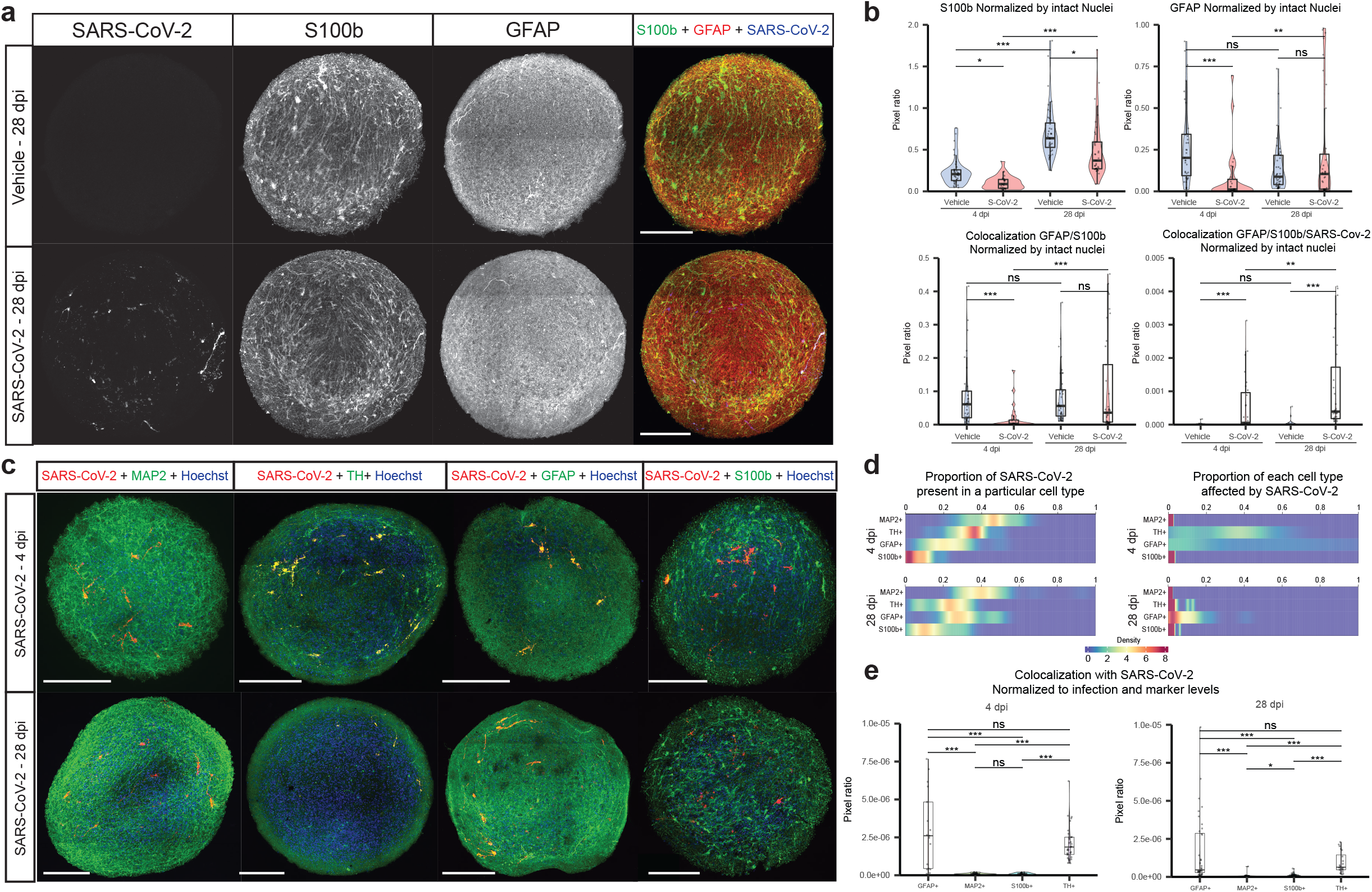
Dopaminergic neurons are highly susceptible to SARS-CoV-2 infection. **a**, Representative images of midbrain organoid sections stained for SARS-CoV-2 Nucleocapsid (N), Glial Fibrillary Acidic Protein (GFAP) and S100 Calcium Binding Protein B (S100b) for the long-term culture (28 dpi) and mock condition. Separate channel images are presented in grey scale, combined images with GFAP in red, S100b in green and SARS-CoV-2 NC in blue. Scale bar, 200μm. **b**, Quantification of pixels of the respective masks normalized to a specific marker with the different conditions, Vehicle (V) or SARS-CoV-2 infection (S), at different timepoints, 4 dpi or 28 dpi; *n(V*4/S4/V28/S28): 35/27/40/38 sections, from 4 organoids per group. Top-left panel, represents the pixel ratio between S100b and intact nuclei. Kruskal-Wallis (*P*<0.0001) followed by Dunn’s test adjusted by Benjamini-Hochberg. Top-right panel, represents the pixel ratio between GFAP and intact nuclei. Kruskal-Wallis (*P*<0.0001) followed by Dunn’s test adjusted by Benjamini-Hochberg. Bottom-left panel, represents the pixel ratio between the colocalization of GFAP and S100b, and intact nuclei. Kruskal-Wallis (*P*<0.0001) followed by Dunn’s test adjusted by Benjamini-Hochberg. Bottom-right panel, represents the pixel ratio between the colocalization of GFAP, S100b and SARS-CoV-2 N signal, and intact nuclei. Kruskal-Wallis (*P*<0.0001) followed by Dunn’s test adjusted by Benjamini-Hochberg. **c**, Representative images of midbrain organoids sections stained for SARS-CoV-2 N (red), nuclei (blue) and one marker (green): Microtubule Associated Protein 2 (MAP2), Tyrosine Hydroxylase (TH), GFAP or S100b. Top and bottom panel represent for short- and long-term culture (4 dpi and 28 dpi) respectively. Scale bar in main panel, 200μm. **d**, Densitograms showing the affinity of SARS-CoV-2 for different cell types depicted by the different markers MAP2, TH, GFAP or S100b, at different times post-infection (4 and 28 dpi). Left panel shows the proportion of SARS-CoV-2 positive pixels colocalizing with one of these markers. Right panel shows the proportion of one of these markers that colocalizes with SARS-CoV-2 positive pixels; *n*(4-MAP2/4-TH/4-GFAP/4-S100b/28-MAP2/28-TH/28-GFAP/28-S100b) 27/33/21/21/41/20/37/37 sections, from 4 organoids per group. **e**, Quantification of SARS-CoV-2 positive pixels colocalized with the different markers MAP2, TH, GFAP or S100b normalized to SARS-CoV-2 positive pixels and the respective marker total quantification. Left panel shows short-culture post-infection (4 dpi), and right panel shows long-culture post-infection (28 dpi). *n*(4-MAP2/4-TH/4-GFAP/4-S100b/28-MAP2/28-TH/28-GFAP/28-S100b) 27/33/21/21/41/20/37/37 sections, from 4 organoids per group. Kruskal-Wallis (*P*<0.0001 for both 4 and 28 dpi) followed by Dunn’s test adjusted by Benjamini-Hochberg. Individual data points are shown with their distribution, and boxplot representing the 25^th^ (lower hinge), median (thick line), and 75^th^ quartile. Whiskers represent 1.5*IQR (inter-quartile range). \**P*<0.05, \*\**P*<0.01, \*\*\**P*<0.001, and ns (not significant) *P*>0.05.

Differentially expressed genes (DEGs) of midbrain organoids 4 dpi were enriched using different bioinformatic platforms. Dysregulated pathways associated with DNA damage, cell stress and death, neurodevelopment and neuronal survival, vesicle transport and membrane recycling, COVID-19, and autophagy were enriched using a manually curated disease centric data base (Fig. 3a). These pathways revealed DEGs such as *RAB7A*, *CTSL*, *VPS26A*, *VPS29*, *VPS35*, *COMMD2*, *PPID*, and *ATP6V1E1* (Extended Data Fig. 2, and Supplementary Table 1 and 2), which were previously reported to be altered post-infection with SARS-CoV-2^15^. Networks of DEGs per category were built to identify the main interactions between them (Fig. 3b and Extended Data Fig. 3). Our results show that the virus is sequestering the machinery of the cell for viral replication by upregulating genes related to its translation (Fig. 3b). This hijacking process involves the endosomal pathway and induces alterations to the dynein axonal transport (Fig. 3b). Concomitant with infection, pathways related to DNA damage, cell stress and apoptosis were also triggered (Fig. 3a and Extended Data Fig. 3). Genes known to be involved in dopaminergic neuronal migration such as *ROBO4* and *SLIT2*^16^, and genes regulating survival of mature neurons like *NOTCH1*^17^ were downregulated post-infection (Fig. 3b), in line with the observed TH+ neuronal fragmentation and cell death. Using other gene sets to enrich the DEGs, pathways related to PD, oxidative phosphorylation, and antigen presentation via MHC class I were activated by SARS-CoV-2 (Fig. 3c,e and Extended Data Fig. 4), while another disease-based gene set showed that annotations related with mitochondrial diseases and movement disorders were significantly dysregulated post-infection (Extended Data Fig. 5). Upregulation of exogenous peptide presentation (Fig. 3d) together with a reduction of “don’t-eat-me” signals (Extended Data Fig. 5), as previously reported in SARS-CoV-2 infected neurons^10^, shows that the infected cell prepares to be cleared. Given that around 20% (921/4438) of the DEGs were non-coding genes (Supplementary Table 3), we analyzed the genomic context of their loci to identify regulatory domains that could be altered by infection^18, 19^. Indeed, genomic regions related to abnormal nervous system physiology and morphology, and behavioral abnormality matched those of the dysregulated non-coding genes (Fig. 3e and Extended Data Fig. 6), with one set of genes related with abnormal morphology of the midbrain by MRI (HP:0002419 - Molar tooth sign on MRI, Fig. 3e). Altogether, this data demonstrates that SARS-CoV-2 infection of midbrain organoids triggers known mechanisms for its replication while inducing cellular damage that can lead to neurodegeneration.

**Fig. 3.**
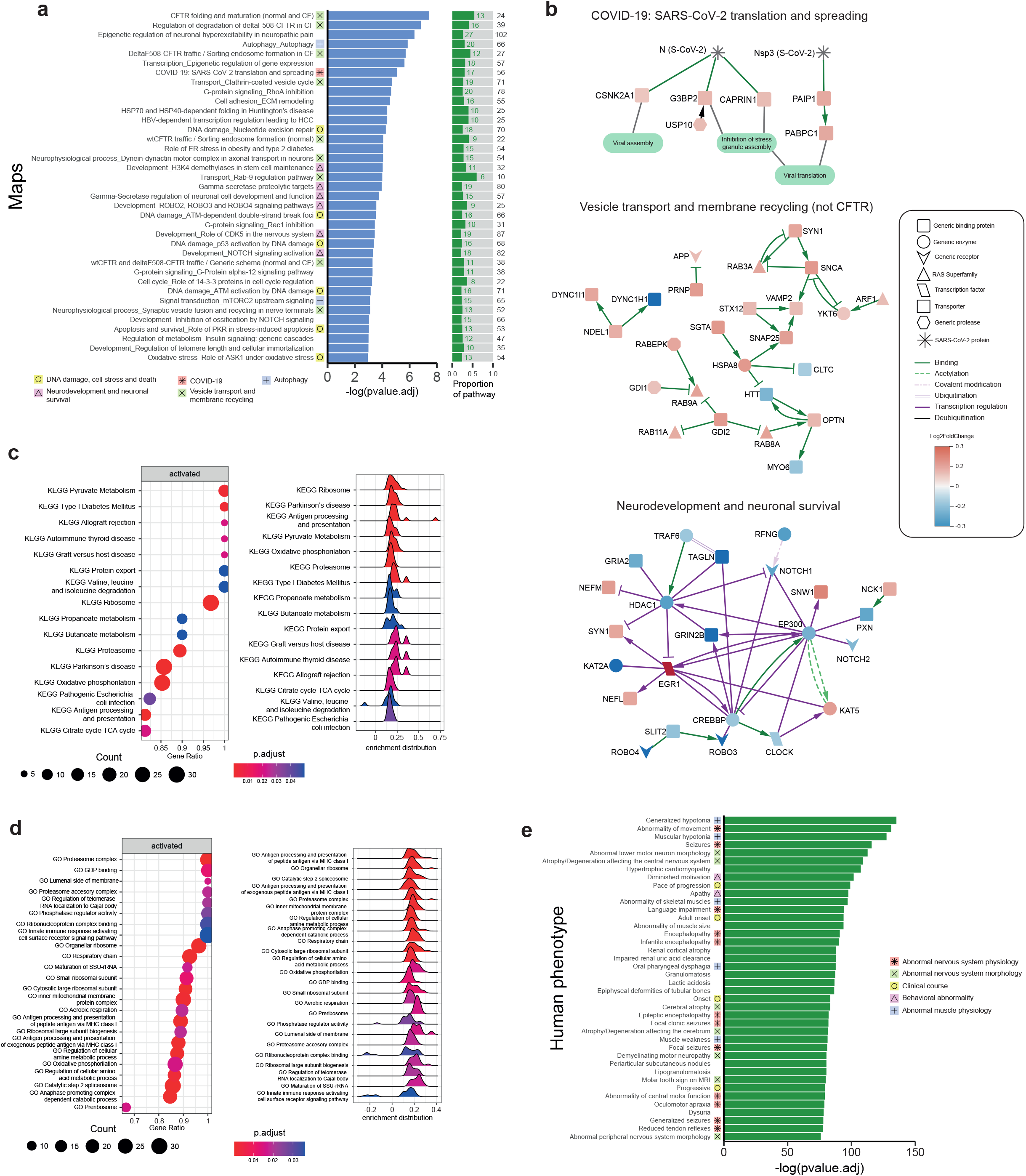
SARS-CoV-2 induces alterations in pathways related to neuronal metabolism and survival. **a**, Significantly dysregulated pathways (*P*<0.05) of DEGs enriched using Metacore gene set with a False discovery rate (FDR) q<0.05. Dysregulated pathways are divided in groups based on their similarity. The proportion of dysregulated genes in the pathway are shown in the right panel (DEGs in green, total genes in black). **b**, Network analysis of DEGs categorized in groups based on the similarity of the dysregulated pathways (grouping shown in panel a). Groups displayed are: COVID19 (C), Vesicle transport and membrane recycling excluding interactions with *CFTR* (VT), and neurodevelopment and neuronal survival (NDS). Networks were filtered to remove isolated nodes, and based on their log2 fold change (log2FC): C and VT < −0.12 or >0.09; NDS < −0.135 or >0.153. **c**, Significantly dysregulated pathways (*P*<0.05) of DEGs enriched using KEGG gene set, with a FDR q<0.05. Dot plot panel, shows the number of DEGs of a specific dysregulated pathway (size of circle), the proportion of dysregulated genes in that pathway (value of Gene ratio), and the adjusted p-value (color of circle). Ridge plot panel, shows the expression (log2FC) distribution of the DEGs of a specific pathway; the color of the histogram represents the adjusted p-value. **d**, Significantly dysregulated pathways (*P*<0.05) of DEGs enriched using GO gene set, with a FDR q<0.05. Dot plot panel, shows the number of DEGs of a specific dysregulated pathway (size of circle), the proportion of dysregulated genes in that pathway (value of Gene ratio), and the adjusted p-value (color of circle). Ridge plot panel, shows the expression (log2FC) distribution of the DEGs of a specific pathway; the color of the histogram represents the adjusted p-value. **e**, Phenotypic abnormalities linked to the loci of non-coding RNA (ncRNA) DEGs, enriched using GREAT, and the Human Phenotype Ontology (HPO) gene set. Dysregulated pathways are divided in groups based on the hierarchy of HPO. Significantly dysregulated pathways by region-based binomial (*P*<0.05), with a FDR q<0.05. RNAseq runs were performed pooling 5 organoids per line and per batch. Vehicle (V) or SARS-CoV-2 infection (S); *n*(V/S): 8/8 RNAseq runs.

Despite several studies have addressed the impact of SARS-CoV-2 in brain organoids^9, 10^, there was a lack of evidence of the effects of viral infection in midbrain organoids at different culture stages. Here we confirm that SARS-CoV-2 has the ability to infect dopaminergic neurons, and that this infection triggers a series of mechanisms that lead to neurite fragmentation and neuronal loss. Furthermore, SARS-CoV-2 induced significant changes in the transcriptome. Pathways centered in membrane recycling, which play a major role in neurons by recycling synaptic vesicles^20^, were among the most dysregulated. SARS-CoV-2 infection also induced dysregulation of dynein-mediated axonal transport, which coincides with previous knowledge that its impairment leads to neuronal death due to the lack of positive feedback from target-derived neurons towards the neuronal soma^21, 22^. Due to their ramified arborization, dopaminergic neurons have a higher susceptibility to altered bi-directional vesicle transport, and such alterations have been linked to the early stages of PD development^23, 24^. Competition for the proteins involved in vesicle recycling seems to be one of the mechanisms underlying SARS-CoV-2-related dopaminergic neuron loss. In addition, these high energy demanding neurons are further affected by impairments in mitochondria metabolism^25^. Our observations are consistent with the previously reported neurotropism of the virus^9^, and highlight the need for further studies to evaluate the interplay between dopaminergic neurons, the blood-brain barrier, and microglia at late infection stages. Our findings expand the knowledge about the neurodegenerative process that the SARS-CoV-2 virus can induce, especially to dopaminergic neurons, emphasizing the importance of understanding the mechanisms underlying long-term neurological impairment in COVID-19 patients.

## Notes

### Competing Interest Statement

J.J. and J.C.S. are co-founders of OrganoTherapeutics SARL.

## References

1. Hu, B., Guo, H., Zhou, P. & Shi, Z.-L. Characteristics of SARS-CoV-2 and COVID-19. Nature Reviews Microbiology 19, 141–154 (2021).

2. Mantovani, A., et al. Long Covid: where we stand and challenges ahead. Cell Death & Differentiation 29, 1891–1900 (2022).

3. Stein, S.R., et al. SARS-CoV-2 infection and persistence in the human body and brain at autopsy. Nature 612, 758–763 (2022).

4. Matschke, J., et al. Neuropathology of patients with COVID-19 in Germany: a post-mortem case series. The Lancet Neurology 19, 919–929 (2020).

5. Zhang, P.-P., et al. COVID-19-associated monocytic encephalitis (CAME): histological and proteomic evidence from autopsy. Signal Transduction and Targeted Therapy 8, 24 (2023).

6. Douaud, G., et al. SARS-CoV-2 is associated with changes in brain structure in UK Biobank. Nature 604, 697–707 (2022).

7. Yong, S.J. Persistent Brainstem Dysfunction in Long-COVID: A Hypothesis. ACS Chemical Neuroscience 12, 573–580 (2021).

8. Han, Y., Yang, L., Lacko, L.A. & Chen, S. Human organoid models to study SARS-CoV-2 infection. Nature Methods 19, 418–428 (2022).

9. Ramani, A., Pranty, A.-I. & Gopalakrishnan, J. Neurotropic Effects of SARS-CoV-2 Modeled by the Human Brain Organoids. Stem Cell Reports 16, 373–384 (2021).

10. Samudyata, et al. SARS-CoV-2 promotes microglial synapse elimination in human brain organoids. Molecular Psychiatry 27, 3939–3950 (2022).

11. Monzel, A.S., et al. Derivation of Human Midbrain-Specific Organoids from Neuroepithelial Stem Cells. Stem Cell Reports 8, 1144–1154 (2017).

12. Boussaad, I., et al. A patient-based model of RNA mis-splicing uncovers treatment targets in Parkinson’s disease. Science Translational Medicine 12, eaau3960 (2020).

13. Jarazo, J., et al. Parkinson’s Disease Phenotypes in Patient Neuronal Cultures and Brain Organoids Improved by 2-Hydroxypropyl-ß-Cyclodextrin Treatment. Movement Disorders 37, 80–94 (2022).

14. Escartin, C., et al. Reactive astrocyte nomenclature, definitions, and future directions. Nature Neuroscience 24, 312–325 (2021).

15. Daniloski, Z., et al. Identification of Required Host Factors for SARS-CoV-2 Infection in Human Cells. Cell 184, 92–105.e116 (2021).

16. Dugan, J.P., Stratton, A., Riley, H.P., Farmer, W.T. & Mastick, G.S. Midbrain dopaminergic axons are guided longitudinally through the diencephalon by Slit/Robo signals. Molecular and Cellular Neuroscience 46, 347–356 (2011).

17. Ables, J.L., Breunig, J.J., Eisch, A.J. & Rakic, P. Not(ch) just development: Notch signalling in the adult brain. Nature Reviews Neuroscience 12, 269–283 (2011).

18. McLean, C.Y., et al. GREAT improves functional interpretation of cis-regulatory regions. Nature Biotechnology 28, 495–501 (2010).

19. Marston, J.L., et al. SARS-CoV-2 infection mediates differential expression of human endogenous retroviruses and long interspersed nuclear elements. JCI Insight 6, e147170 (2021).

20. Watanabe, S., et al. Clathrin regenerates synaptic vesicles from endosomes. Nature 515, 228–233 (2014).

21. Heerssen, H.M., Pazyra, M.F. & Segal, R.A. Dynein motors transport activated Trks to promote survival of target-dependent neurons. Nature Neuroscience 7, 596–604 (2004).

22. LaMonte, B.H., et al. Disruption of Dynein/Dynactin Inhibits Axonal Transport in Motor Neurons Causing Late-Onset Progressive Degeneration. Neuron 34, 715–727 (2002).

23. Abeliovich, A. & Gitler, A.D. Defects in trafficking bridge Parkinson’s disease pathology and genetics. Nature 539, 207–216 (2016).

24. Bellucci, A., Longhena, F. & Spillantini, M.G. The Role of Rab Proteins in Parkinson’s Disease Synaptopathy. Biomedicines 10, 1941 (2022).

25. Fu, H., Hardy, J. & Duff, K.E. Selective vulnerability in neurodegenerative diseases. Nature Neuroscience 21, 1350–1358 (2018).

