## Supplementary Information for "SARS-CoV-2 infection induces dopaminergic neuronal loss in midbrain organoids during short and prolonged cultures"

**Materials and Methods**

**Information about cells used in the manuscript**

Two healthy human induced pluripotent stem cell (hiPSC) control lines were used in all experiments. These lines were acquired from Alstem (https://www.alstembio.com). Their reference number is iPS11 and iPS15. These lines were derived from foreskin fibroblast and peripheral blood mononuclear cells respectively, and reprogrammed by episomal plasmids. The VeroE6 cells used for viral isolation were a kind gift from Dr. Thorsten Wolff, Influenza und respiratorische Viren, Robert Koch Institute, Germany.

**hiPSC culture, and NESCs generation**

hiPSCs were cultured with Essential 8 (E8) medium (Thermo Fisher Scientific, A1517001) in Matrigel (Corning, 354277) coated 6-well plates under 5%CO_2_. Cultures were passaged either with Accutase (Sigma, A6964) when a single cell preparation was needed or 1:1,000 EDTA (Thermo Fisher Scientific, 15575020) in DPBS (Thermo Fisher Scientific, 14190250) for expansion. Cells were cultured with E8 supplemented with 10 μM ROCK inhibitor Y-27632 (Merck, 688000) overnight. Neuroepithelial stem cells (NESCs, the starting point used for organoid derivation) were generated by making embryoid bodies (EBs) in an AggreWellTM400 plate (STEMCELL Technologies, 27845). hiPSCs were cultured until a confluency of ~80% determined by visual inspection under an inverted microscope (Nikon Eclipse TS100-F Eclipse, RRID:SCR_020324) was reached. The culture medium was aspirated and the cells were treated with 1 ml of Accutase per well for 5 minutes in the incubator. Detached cells were further pipetted to single cells, collected in a 15 ml centrifuge tube with 10ml of E8 and centrifuge at 300*g* for 3 min. Cells were then resuspended and plated in an AggreWell plate using EBs generation medium consisting of 1x KO-DMEM (Thermo Fisher Scientific, 10829018), 1:5 Knockout serum replacement (Thermo Fisher Scientific, 10828028), 1:1,000 Penicillin-Streptomycin (Thermo Fisher Scientific, 15140122), 1:1,000 GlutaMAX (Thermo Fisher Scientific, 35050038) and 1:1,000 NEAA (Thermo Fisher Scientific, 11140050), and 1:5,000 β-mercaptoethanol (Thermo Fisher Scientific, 31350010) supplemented with 3 μM	CHIR 99021 (Axon Medchem, CT99021), 10 μM SB‐431542 (Abcam, ab120163), 1 μM dorsomorphin (Tocris, 3093), 0.75 μM puromorphamine (Enzo Life Science, ALX-420-045), and 10 μM ROCK inhibitor Y-27632. The AggreWell plate was prepared following the manufacturer’s instructions. 1.2 million cells (counted with Life Technologies Countess Automated Cell Counter, RRID:SCR_020236, following manufacturer’s protocol) were plated and the AggreWell plate was centrifuged at 100*g* for 3min. This stage is considered the start of the culture (day 0) of the NESCs generation. After 24 hours, the formation of EBs was assessed under an inverted microscope, and EBs were collected by gently pipetting medium up and down to let all the EBs float. The EBs’ suspension was transferred into a 15 ml tube and let to sediment for 2 min. Medium was aspirated and EBs resuspended in 5 ml of EB supplemented medium (without ROCK inhibitor) and transferred to a low attachment tissue culture 6-well plate (Corning, 3471). EBs were cultured for 5 days more, changing the media to N2B27 patterning and N2B27 maintenance medium on days 2 and 4 from the start of the culture respectively. Both patterning media and maintenance medium have the same basal composition (known as N2B27) but are supplemented differently. N2B27 medium consists of equal quantities of DMEM/F12 (Thermo Fisher Scientific, 21331020) and Neurobasal (Thermo Fisher Scientific, 21103049), with 1:1,000 Penicillin-Streptomycin (Thermo Fisher Scientific, 15140122), 1:1,000 GlutaMAX (Thermo Fisher Scientific, 35050038), 1:1,000 B27 without vitamin A (Thermo Fisher Scientific, 12587010), and 1:2,000 N2 Supplement (Thermo Fisher Scientific, 17502048). The patterning medium N2B27 is supplemented with the same supplements as EB generation media (without ROCK inhibitor) while the maintenance media N2B27 is supplemented with 3 μM CHIR 99021 (Axon Medchem, CT99021), 0.75 μM puromorphamine (Enzo Life Science, ALX-420-045), and 150 μM ascorbic acid (Merck, A4544). On day 6, EBs forming neural-tube like structures were transferred to a 15ml tube to sediment for 2 min. Using a pipette, EBs were shredded by pipetting up and down 5 to 10 times, until pieces are barely visible by eye. The cell suspension was plated in Matrigel-coated 12-well plates. Purification of the newly generated NESCs was performed by passaging the colonies via dissociation with Accutase incubated for 5 min in the incubator, diluting the Accutase with 10ml of N2B27 in a 15ml tube, centrifuging at 300*g* for 3 min, and seeding 72,000 cells/cm^2^. Normally, 5 passages (once or twice per week) are needed to purify the NESCs.

**Midbrain organoid generation and culture**

Cultured NESCs were detached as described in the previous section, and 9000 cells/well were plated in 100μl of N2B27 maintenance medium in a 96-well ultra-low adhesion plate (Corning, 7007). The plate was centrifuged at 300g for 3 min, incubated in 5%CO_2_, and media was changed every 2 days. After 4 days, medium was changed to differentiation medium (considering this day 0 of the organoids) consisting of N2B27 supplemented with 10 ng/ml hBDNF (Peprotech, 450-02), 10 ng/ml hGDNF (Peprotech, 450-10), 500 μM db cAMP (Merck, D0627), 1ng/ml TGF-β3 (Peprotech, 100-36E), 1 μM puromorphamine (Enzo Life Science, ALX-420-045), and 150 μM ascorbic acid (Merck, A4544). Medium was changed every other day. On day 7 of differentiation, medium was switched to maturation medium (puromorphamine removed from the supplements). Organoids were cultured for 23-26 days more in maturation medium (30-33 days old organoids). Four batches of organoids were generated in successive days (8 organoids per batch per hiPSC line), and transferred together to a BSL-3 lab to perform the infection.

**Viral infection**

SARS-CoV-2 B.1 viral strain was isolated from a patient with confirmed COVID-19 in Luxembourg during the first wave of the pandemic (before April 15th 2020). This strain harbors the D614G mutation as well as a 42-amino acid deletion (nucleotides 28040 and 28165) in the ORF8 gene (European Nucleotide Archive, PRJEB59248, https://www.ebi.ac.uk/ena/browser/home). It was isolated and titrated using the Reed and Muench method^1^ on VeroE6 cells. Sequencing was carried out to confirm sequence integrity. Whole Genome Sequencing was carried out on the Illumina iSeq100 upon viral RNA extraction. The Spike protein sequence was extracted from the fasta files and aligned to the Wuhan-1 reference spike protein. Midbrain organoids were exposed to 0.05 MOI of SARS-CoV-2 virus overnight at 37°C, then washed twice with PBS and incubated in differentiation medium for 4- or 28-days post-infection (dpi). For cultures exceeding 4 dpi, medium was replaced weekly.

**Immunofluorescence**

Three organoids per batch per cell line were fixed by incubating the organoids in 4% formaldehyde (Merck, 1.00496) overnight. The organoids were washed 3 times with DPBS, and embedded in 3% low melting point agarose (Biozym, 840101) in water, incubated for 15 min at 37 °C, and kept 30 min at room temperature. The solidified agarose block was covered in DPBS and kept overnight at 4 °C before sectioning. Sections were generated with a Vibratome VT1000S (Leica Microsystems, RRID:SCR_016495) of a thickness of 50 µm at a speed of 6 and a frequency of 8. Sections coming from the same organoid were pooled and transferred as free floating sections to a well of a 24-well plate where the staining protocol was performed. Sections were incubated in 2% normal goat serum (Thermo Fisher Scientiﬁc, 10000C), 2% bovine serum albumin (Merck, A4503) and 0.5% Triton X-100 (Carl Roth, 3051.3) in DPBS (Blocking solution) for 90 min on a shaker at room temperature. Staining was performed for 48h in Blocking solution containing 0.1% Triton X-100 and the primary antibodies on a shaker at 4 °C. The antibody combinations for Fig. 1, Fig. 2c, and Extended data Fig. 1a included chicken anti-Tyrosine Hydroxylase (Abcam, ab76442, RRID:AB_1524535) at 1:500, and rabbit anti-SARS-CoV-2 Nucleocapsid (Genetex, GTX635679, RRID:AB_2888553) at 1:1,000. The antibody combination for Extended data Fig. 1b included mouse anti- SARS-CoV-2 nucleocapsid (Sino Biological, 40143-MM05, RRID:AB_2827977) at 1:1,000, and rabbit anti-Tyrosine Hydroxylase (Abcam, ab112, RRID:AB_297840), at 1:500. The antibody combinations for Fig. 2a and c included chicken anti-GFAP (Merck, AB5541, RRID:AB_177521) at 1:1,000, mouse anti-S-100 beta-subunit (Merck, S2532, RRID:AB_477499) at 1:1,000, and rabbit anti-SARS-CoV-2 Nucleocapsid (Genetex, GTX635679, RRID:AB_2888553) at 1:1,000. The antibody combination for Fig. 2c included mouse anti- SARS-CoV-2 Nucleocapsid (Sino Biological, 40143-MM05, RRID:AB_2827977) at 1:1,000, and chicken anti-MAP2 (Abcam, ab92434, RRID:AB_2138147). After incubation with the primary antibody solution, sections were washed 3 times with 500 µl of DPBS and incubated with the secondary antibody solution consisting of 2% normal goat serum, 2% bovine serum albumin, and 0.05 % Tween-20 (Merck, P7949) in DPBS for 2 hours at room temperature. Nuclear staining with the dye Hoechst 33342 (Thermo Fisher Scientiﬁc, H21492) at 1:1,000 was performed at the time of incubation with the secondary antibodies. The secondary antibodies used were goat anti-mouse IgG 488 (Thermo Fisher Scientiﬁc, A-11001, RRID:AB_2534069), goat anti-rabbit IgG 568 (Thermo Fisher Scientiﬁc, A-11036, RRID:AB_10563566), and goat anti-chicken IgG 647 (Thermo Fisher Scientiﬁc, A-21449, RRID:AB_2535866). All secondary antibodies were used at 1:1,000 dilution. Sections were mounted under a stereomicroscope (Carl Zeiss Stemi DV4 Stereomicroscope, RRID:SCR_021244) on 30-well Teflon coated slides (De Beer Medicals, BM-9133) with Fluoromount-G (SouthernBiotech, 0100-01) and covered with a 24x50mm cover glass (VWR, 631-1574). Slides were imaged after 24 hours.

**Image acquisition**

Imaging of the organoids’ sections was performed with fully automated microscope Yokogawa CV8000 using the SearchFirst modality of the Wako Software with a Matlab script (see section Code availability), followed by a second pass with a grid pattern since hits were not centered. Script parameters of intensity higher than 4000, area filter for structures between 2000 and 5e6 pixels, and a morphological close with the function imclose of the segmented image with a disk structural element of 51 were used to automatically segment the sections at 4x. The function bwlabeln was used for labelling the identified structures. Their position was identified by creating a point grid to match with the localization of the segmented structure, and re-scanned at 20x air objective for acquisition of the final images. The first pass was used to recognize the nuclear staining (Hoechst) with a 4x objective, a light source of 405nm (power of 80%), a band pass filter of 445/45nm, an exposure time of 150ms, binning 4 and a pinhole diameter of 50µm. The second pass settings were 20x air objective, light sources with their respective power and band pass filter of 405nm (power of 80%,445/45nm), 488nm (power of 50%, 525/50nm), 561nm (power of 80%, 600/37nm) and 640nm (power of 80%, 676/29nm), all with exposure times of 50ms, binning 1, and a pinhole diameter of 50µm. A Z-stack of 21 planes with a distance of 2µm between them was acquired for all channels.

**Image analysis**

Images acquired in the second pass were analyzed using a Matlab (The MathWorks, R2021a) script (see section Code availability). Loaded images were stitched and the planes with best signal were automatically selected by combining all the channels and filtering planes with a signal of at least 66% of the maximum signal. Channels were analyzed separately and masks generated from them were used for detecting colocalization between the channels^2^. Nuclei segmentation was achieved by neighborhood processing the pixels (to increase the difference with the background) with a difference of Gaussian, in the spatial domain. The foreground image was convoluted with a Gaussian filter size (GFS) of 101 pixels and a standard deviation (SD) of 3 pixels. The background image to subtract was convoluted with a 101 pixels GFS and a SD of 7 pixels. Connected elements that had an intensity higher than 50 and a size bigger than 300 pixels were included in the nuclei mask. To identify pyknotic nuclei a convolution with a GFS of 11 pixels and a SD of 1 pixel was performed and pixels with a value higher than 4000 were included in the pyknotic mask. Segmentation of the non-nuclear channels was performed in the frequency domain by performing a Fourier transform per plane followed by an inverse Fourier transform to return the images to the spatial domain. A Butterworth high pass filtering was used in the frequency domain to sharpen the images with cutoff frequency of 7 or 15, and an order of 1. Parameters for the thresholds for constructing the final mask were chosen by visual inspection of representative images of each channel, and applied to all the image set per channel. Identification of neurite fragmentation was performed by erosion with a sphere structural element. Representative images were generated using the imadjust function with fixed parameters per channel across all images.

**RNAseq and differentially expressed gene analysis**

For gene expression analyses, 5 organoids per line and per batch were pooled, and RNA was extracted using the QIAshredder (Qiagen, 79654) and the RNeasy Mini kit (Qiagen, 74104) following manufacturer’s protocol after 4 dpi. A total of 750 ng of RNA was used for library preparation using TruSeq stranded mRNA library preparation kit (Illumina, 20020594) following the provided protocol. Briefly, mRNA pull down was done using magnetic beads with oligodT primer. To preserve strand information, the second strand synthesis was done such that during PCR amplification only the first strand was amplified. The libraries were quantified using the Qubit dsDNA HS assay kit (Thermo Fisher Scientific, Q32851) and the size distribution was determined using an Agilent 2100 Bioanalyzer (RRID:SCR_019389). Pooled libraries were sequenced at University of Luxembourg LCSB Sequencing Platform Core Facility (RRID:SCR_021931) using NextSeq500. The raw RNA sequencing data was pre-processed using the software package Rsubread^3^ of R^4^. Gene-level differential expression analysis was conducted in R using the software package DESeq2^5^ and filtering out genes with low expression counts (sum of counts < 10 across all samples).

**Gene enrichment and network analysis**

Pathway enrichment and network analyses were implemented in the GeneGo MetaCore™ software (https://portal.genego.com) using the gene-level differential expression analysis results obtained with DESeq2 as input for the standard enrichment analysis workflow. The pathway over-representation analysis statistics, including false-discovery rate (FDR) scores according to the method by Benjamini and Hochberg^6^, were determined for the GeneGo collections of cellular pathway maps and process networks. Network visualizations were created using the standard workflow of the "Build Network for Single Gene/Protein/Compound or a List" function in GeneGo, and filtering the network data to cover only the human species and to exclude the object types "Compound", "RNA", "DNA", "Predicted metabolite", and "Inorganic ion" from the analysis. Categories of the dysregulated pathways coming from MetaCore™ were established based on their similarity. Enrichments with Gene Ontology (GO)^7, 8^, Kyoto Encyclopedia of Genes and Genomes (KEGG)^9, 10^, and Reactome^11, 12^ gene sets were done using the GSEA function from the clusterProfiler^13^ package of R, with a minimal size of genes annotated for testing of 5 and maximal gene set for analyzing of 500, a *P* value cutoff of 0.05 and FDR as adjustment method. The molecular signatures database (MSigDB) used for all the gene sets was v6.2. Enrichments with the DisGeNET (DGN)^14^ and Disease ontology (DO)^15^ gene set were performed with the enrichDGN and enrichDO functions from the DOSE^16^ package of R with default parameters. Networks were constructed based on the similarity of the dysregulated pathways obtain from MetaCore™ (groups displayed in Fig. 3a). Nodes and interactions files were generated with MetaCore™. Networks were filtered to remove isolated nodes, and based on the log2 fold change using Cytoscape^17^ (<https://cytoscape.org/>).

**Genomic region enrichment**

Determination of the dysregulated genomic regions using the non-coding RNAs (ncRNAs) detected in the DEGs was performed using the GREAT^18, 19^ software v4.0.4 with default settings, and significance by region-based binomial. ncRNAs from the analysis with Metacore (in GeneSymbol format) were converted to UCSC ID with bioDBnet^20^ (<https://biodbnet-abcc.ncifcrf.gov>), and further converted to Browser Extensible Data (BED) format with the UCSC Table browser^21^ (<http://genome.ucsc.edu/cgi-bin/hgTables>).

**Statistical analyses**

Statistical analyses as well as the exact *n* are mentioned in each figure legend. For each of the immunofluorescence staining experiments, two organoids from each cell line per condition were sectioned, and all the recovered sections were stained, imaged and automatically analyzed. For RNAseq, five organoids per cell line, batch, and condition were pooled. Each of these pools was sequenced and analyzed separately. Pooling both cell lines per batch was done to compare the conditions in the imaging and RNAseq analyses. Differentially expressed genes (DEGs) *P* value significance scores were computed using the Wald test, and were adjusted for multiple hypothesis testing according to the Benjamini and Hochberg method. All statistical analyses for the imaging experiments were done in R. Non-parametric tests (Kruskal-Wallis test followed by Dunn’s test with a Benjamini-Hochberg adjustment for multiple comparisons) were used to compare the results of image analyses. All the *P* values reported are two-tailed. Gene enrichments with GO, KEGG, Reactome and DGN were performed in R with the parameters mentioned in the Gene enrichment section. Clustering analysis was performed using the complexheatmap^22^ package of R, with Canberra distance and a complete hierarchical clustering. No statistical method was used to determine the sample size. Sample size was determined based on previous observations within our lab^23^. No blinding of condition was performed for the analysis; an automated analysis of the images and RNAseq data was performed using the available scripts (see Code availability section) with equal settings across conditions. No outlier removal procedure was done, and no repeated measures on the same sample were performed. Individual data points are shown with their distribution (violin plot), and boxplot representing the 25th (lower hinge), median (thick line), and 75th quartile. Whiskers represent 1.5*IQR (inter-quartile range). **P*<0.05, ***P*<0.01, ****P*<0.001, and ns (not significant) *P*>0.05.

**Figure production**

Panels of immunofluorescence representative images with their zoom regions were edited with Illustrator CC (Abode Systems, v27.2). No further image adjustments were made to the immunofluorescence staining after using the imadjust function with fixed parameters across conditions in Matlab. Violin plots, boxplots, volcano plots and bar graphs were produced in R^4, 24-34^. Heatmap and densitograms were produced using the complexheatmap^22^ and circlize^35^ packages of R. Pathway enrichment dot plots and ridge plots were produced with the enrichplot package of R^13^. Display of constructed networks with Cytoscape^17^.

**Reporting summary**

Further information on research design is available in the Nature Research Reporting Summary linked to this article.

**Data availability**

All processed data can be found here: <https://doi.org/10.17881/64fh-dt90>. Due to the size of the raw microscope images, they are available upon request (a data transfer procedure will be organized accordingly). Raw data of the RNAseq experiments can be accessed at Gene Expression Omnibus under accession number GSE225517.

**Code availability**

All the R and Matlab codes used in this manuscript can be found in <https://gitlab.lcsb.uni.lu/dvb/jarazo-et-al_2023>.

**Acknowledgment**

We thank the LCSB Sequencing Platform Core Facility, and the LCSB Imaging facility for the support in this project. This work was supported by the Ministry of Economy of Luxembourg (20200616RDI170010215073-RDI-REDIND).

**Author contribution**

J.J., D.P.B. and J.C.S. conceptualized and designed the study. J.J. and E.S.S performed the experiments. J.J. performed the image analysis. J.J and E.G. performed the gene expression analyses. J.J. combined and analyzed all the experimental data. J.J. wrote the first version of the manuscript to which all authors contributed with its revision. D.P.B. and J.C.S. supervised.

**Competing interests**

J.J. and J.C.S. are co-founders of OrganoTherapeutics SARL.

**References**

1.	Reed, L.J. & Muench, H. A simple method of estimating fifty per cent endpoints. *American Journal of Epidemiology* **27**, 493-497 (1938).

2.	Marques, O. *Practical Image and Video Processing Using MATLAB®* (John Wiley & Sons, Inc., 2011).

3.	Liao, Y., Smyth, G.K. & Shi, W. The R package Rsubread is easier, faster, cheaper and better for alignment and quantification of RNA sequencing reads. *Nucleic Acids Research* **47**, e47-e47 (2019).

4.	R, C.T. R: A language and environment for statistical computing. (2022).

5.	Love, M.I., Huber, W. & Anders, S. Moderated estimation of fold change and dispersion for RNA-seq data with DESeq2. *Genome Biology* **15**, 550 (2014).

6.	Benjamini, Y. & Hochberg, Y. Controlling the False Discovery Rate: A Practical and Powerful Approach to Multiple Testing. *Journal of the Royal Statistical Society: Series B (Methodological)* **57**, 289-300 (1995).

7.	Ashburner, M.*, et al.* Gene Ontology: tool for the unification of biology. *Nature Genetics* **25**, 25-29 (2000).

8.	The Gene Ontology, C. The Gene Ontology resource: enriching a GOld mine. *Nucleic Acids Research* **49**, D325-D334 (2021).

9.	Kanehisa, M. & Goto, S. KEGG: Kyoto Encyclopedia of Genes and Genomes. *Nucleic Acids Research* **28**, 27-30 (2000).

10.	Kanehisa, M. Toward understanding the origin and evolution of cellular organisms. *Protein Science* **28**, 1947-1951 (2019).

11.	Wu, G. & Haw, R. Functional Interaction Network Construction and Analysis for Disease Discovery. in *Protein Bioinformatics: From Protein Modifications and Networks to Proteomics* (ed. C.H. Wu, C.N. Arighi & K.E. Ross) 235-253 (Springer New York, New York, NY, 2017).

12.	Gillespie, M.*, et al.* The reactome pathway knowledgebase 2022. *Nucleic Acids Research* **50**, D687-D692 (2022).

13.	Yu, G., Wang, L.-G., Han, Y. & He, Q.-Y. clusterProfiler: an R Package for Comparing Biological Themes Among Gene Clusters. *OMICS: A Journal of Integrative Biology* **16**, 284-287 (2012).

14.	Piñero, J.*, et al.* The DisGeNET knowledge platform for disease genomics: 2019 update. *Nucleic Acids Research* **48**, D845-D855 (2020).

15.	Schriml, L.M.*, et al.* Human Disease Ontology 2018 update: classification, content and workflow expansion. *Nucleic Acids Research* **47**, D955-D962 (2019).

16.	Yu, G., Wang, L.-G., Yan, G.-R. & He, Q.-Y. DOSE: an R/Bioconductor package for disease ontology semantic and enrichment analysis. *Bioinformatics* **31**, 608-609 (2015).

17.	Shannon, P.*, et al.* Cytoscape: a software environment for integrated models of biomolecular interaction networks. *Genome Res* **13**, 2498-2504 (2003).

18.	McLean, C.Y.*, et al.* GREAT improves functional interpretation of cis-regulatory regions. *Nature Biotechnology* **28**, 495-501 (2010).

19.	Tanigawa, Y., Dyer, E.S. & Bejerano, G. WhichTF is functionally important in your open chromatin data? *PLOS Computational Biology* **18**, e1010378 (2022).

20.	Mudunuri, U., Che, A., Yi, M. & Stephens, R.M. bioDBnet: the biological database network. *Bioinformatics* **25**, 555-556 (2009).

21.	Karolchik, D.*, et al.* The UCSC Table Browser data retrieval tool. *Nucleic Acids Research* **32**, D493-D496 (2004).

22.	Gu, Z., Eils, R. & Schlesner, M. Complex heatmaps reveal patterns and correlations in multidimensional genomic data. *Bioinformatics* **32**, 2847-2849 (2016).

23.	Brown, S.J.*, et al.* PINK1 deficiency impairs adult neurogenesis of dopaminergic neurons. *Scientific Reports* **11**, 6617 (2021).

24.	Posit, T. RStudio: Integrated Development Environment for R. (2022).

25.	Wickham H, A.M., Bryan J, Chang W, McGowan LD, François R, Grolemund G, Hayes A, Henry L, Hester J, Kuhn M, Pedersen TL, Miller E, Bache SM, Müller K, Ooms J, Robinson D, Seidel DP, Spinu V, Takahashi K, Vaughan D, Wilke C, Woo K, Yutani H. Welcome to the tidyverse. *Journal of Open Source Software* **4**, 1686 (2019).

26.	Ahlmann-Eltze, C. & Pati, I. ggsignif: R Package for Displaying Significance Brackets for 'ggplot2'. *PsyArXiv* (2021).

27.	Kassambara, A. ggpubr: 'ggplot2' Based Publication Ready Plots. (2022).

28.	Wilke, C. cowplot: Streamlined Plot Theme and Plot Annotations for 'ggplot2'. (2020).

29.	Dinno, A. dunn.test: Dunn's Test of Multiple Comparisons Using Rank Sums. (2017).

30.	Dowle, M. & Srinivasan, A. data.table: Extension of `data.frame`. (2022).

31.	Wickham, H. & Bryan, J. readxl: Read Excel Files. (2022).

32.	Wickham, H. & Seidel, D. scales: Scale Functions for Visualization. (2022).

33.	Waldron, L. & Riester, M. HGNChelper: Identify and Correct Invalid HGNC Human Gene Symbols and MGI Mouse Gene Symbols. (2019).

34.	Neuwirth, E. RColorBrewer: ColorBrewer Palettes. (2022).

35.	Gu, Z., Gu, L., Eils, R., Schlesner, M. & Brors, B. circlize implements and enhances circular visualization in R. *Bioinformatics* **30**, 2811-2812 (2014).

**Extended data figure legends**

**Extended data Fig. 1** **| SARS-CoV-2 colocalization with dopaminergic neuron marker TH. a**, Images of a midbrain organoid section (same section as Fig. 1a) stained for SARS-CoV-2 Nucleocapsid (N), Tyrosine Hydroxylase (TH) and a nuclear dye (Hoechst) for the short-term culture (4 dpi) and mock condition. Separate channel images are presented in grey scale, and automated segmentation mask in red. Combined images with SARS-CoV-2 N in red, TH in green and Hoechst in blue, and colocalization mask between SARS-CoV-2 N and TH in white. Boxes represent zoomed regions in the panel below. Scale bar in main panel, 200µm. Scale bar zoomed region, 50 µm. **b**, Representative images of midbrain organoids sections stained for SARS-CoV-2 Nucleocapsid (N), TH and Hoechst for the short-term culture (4 dpi) and mock condition, using different antibodies compared to Fig. 1. Separate channel images are presented in grey scale, combined images with SARS-CoV-2 N in red, TH in green and Hoechst in blue. Boxes represent zoomed regions in the panel below. Scale bar in main panel, 200µm. Scale bar zoomed region, 50 µm.

**Extended data Fig. 2** **| SARS-CoV-2 infection induces dysregulation of vesicle recycling and oxidative phosphorylation. a**, Volcano plot showing the differentially expressed genes (DEGs) between infected and non-infected midbrain organoids at 4 dpi. Genes in grey are not significantly dysregulated (*P*>0.05). Genes in blue passed the fold change (FC) threshold (bigger than 1.5 or smaller than -1.5) but are not significantly dysregulated (*P*>0.05). Genes in green are significantly dysregulated (*P*<0.05) but did not pass the FC threshold. Genes in red are significantly dysregulated (*P*<0.05) and passed the FC threshold. **b**, Heatmap of selected differentially expressed genes (DEGs) grouped based on the dysregulated pathways described in Danilosky et al., 2021. Color and symbol coding represent the enriched pathway the DEGs belong. Canberra distance and a complete hierarchical clustering were used. Each column represents different RNAseq runs pooling 5 organoids each.

**Extended data Fig. 3** **| Network analysis displays the protein interaction of differentially expressed genes altered by SARS-CoV-2 infection. a**, Network analysis of DEGs categorized under the group Autophagy based on the similarity of the dysregulated pathways (grouping shown in Fig. 3a). Network was filtered to remove isolated nodes, and based on their log2 fold change (log2FC) with a threshold of < -0.12 or >0.1. **b**, Network analysis of DEGs categorized under the group DNA damage, cell stress and death based on the similarity of the dysregulated pathways (grouping shown in Fig. 3a). Network was filtered to remove isolated nodes, and based on their log2FC with a threshold of < -0.2 or >0.2. **c**, Network analysis of DEGs categorized under the group Vesicle transport and membrane recycling including only interactions with *CFTR* based on the similarity of the dysregulated pathways (grouping shown in Fig. 3a). Network was filtered to remove isolated nodes, and based on their log2FC with a threshold of < -0.12 or >0.1.

**Extended data Fig. 4** **| Clustering of differentially expressed genes based on dysregulated pathways shows the cellular response induced by SARS-CoV-2. a,** Heatmap of differentially expressed genes (DEGs) grouped based on the top enriched pathways of the KEGG (left) and GO (right) gene sets. Color and symbol coding represent the enriched pathway the DEGs belong. Canberra distance and a complete hierarchical clustering were used. Each column represents different RNAseq runs pooling 5 organoids each. **b**, Significantly dysregulated pathways (*P*<0.05) of DEGs enriched using Reactome gene set, with a False discovery rate (FDR) q<0.05. Dot plot panel, shows the number of DEGs of a specific dysregulated pathway (size of circle), the proportion of genes dysregulated from that pathway (value of Gene ratio), and the adjusted p-value (color of circle). Ridge plot panel, shows the expression (log2FC) distribution of the DEGs of a specific pathway; the color of the histogram represents the adjusted p-value. Differential expression was calculated using the negative binomial distribution of the DESeq2 package from R with a FDR q<0.05. Vehicle (V) or SARS-CoV-2 infection (S); *n*(V/S): 8/8 RNAseq runs. **P*<0.05, ***P*<0.01, ****P*<0.001, and ns (not significant) *P*>0.05.

**Extended data Fig. 5 | Differentially expressed genes after SARS-CoV-2 infection leads to dysregulation of pathways involved in antigen presentation and mitochondrial activity. a**, Significantly dysregulated pathways (*P*<0.05) of DEGs enriched using DisGeNET gene set, with a False discovery rate (FDR) q<0.05. **b**, Significantly dysregulated pathways (*P*<0.05) of DEGs enriched using Disease ontology gene set, with a FDR q<0.05. **c**, Differentially expressed levels of CD46 and CD200. Each dot represents the preprocessed expression count of different RNAseq runs pooling 5 organoids each. Differential expression was calculated using the negative binomial distribution of the DESeq2 package from R with a FDR q<0.05. Vehicle (V) or SARS-CoV-2 infection (S); *n*(V/S): 8/8 RNAseq runs. **P*<0.05, ***P*<0.01, ****P*<0.001, and ns (not significant) *P*>0.05.

**Extended data Fig. 6** **| Dysregulated genomic regions linked to viral replication, mitochondrial metabolism, and microtubule organization and neurite growth. a**, Gene Ontology biological process annotation of significantly dysregulated genomic regions enriched with GREAT, based on differentially expressed non-coding RNA. Significantly dysregulated pathways by region-based binomial (*P*<0.05), with a False discovery rate (FDR) q<0.05. **b**, Gene Ontology cellular component annotation of significantly dysregulated genomic regions enriched with GREAT, based on differentially expressed non-coding RNA. Significantly dysregulated pathways by region-based binomial (*P*<0.05), with a FDR q<0.05. **c**, Gene Ontology molecular function annotation of significantly dysregulated genomic regions enriched with GREAT, based on differentially expressed non-coding RNA. Significantly dysregulated pathways by region-based binomial (*P*<0.05), with a FDR q<0.05.
