## Supplementary figures and images for "SARS-CoV-2 infection induces dopaminergic neuronal loss in midbrain organoids during short and prolonged cultures"

### Supplementary Figure 1

**Exteded data Figure 1**

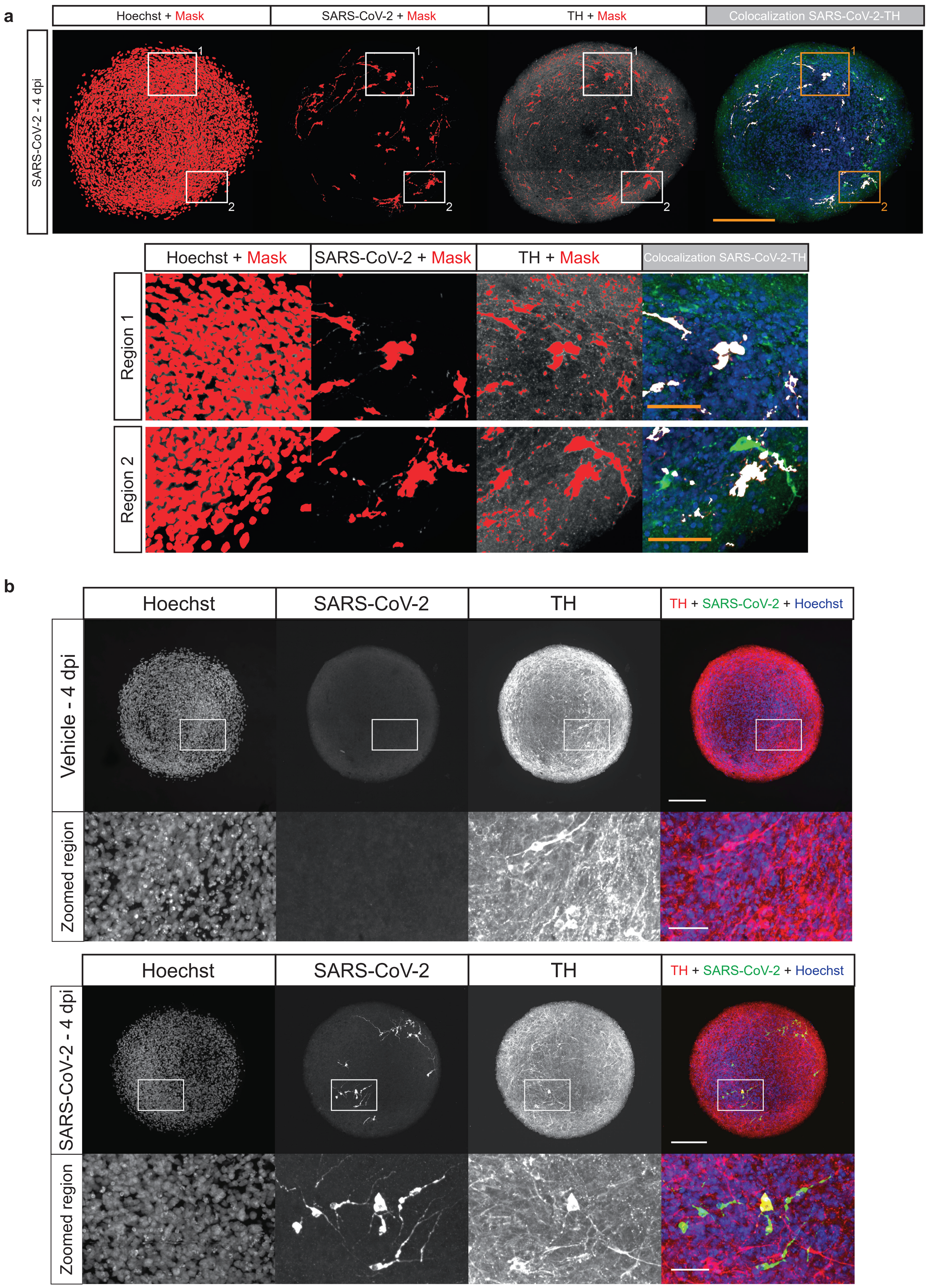

### Supplementary Figure 2

#### Extended data Figure 2

a

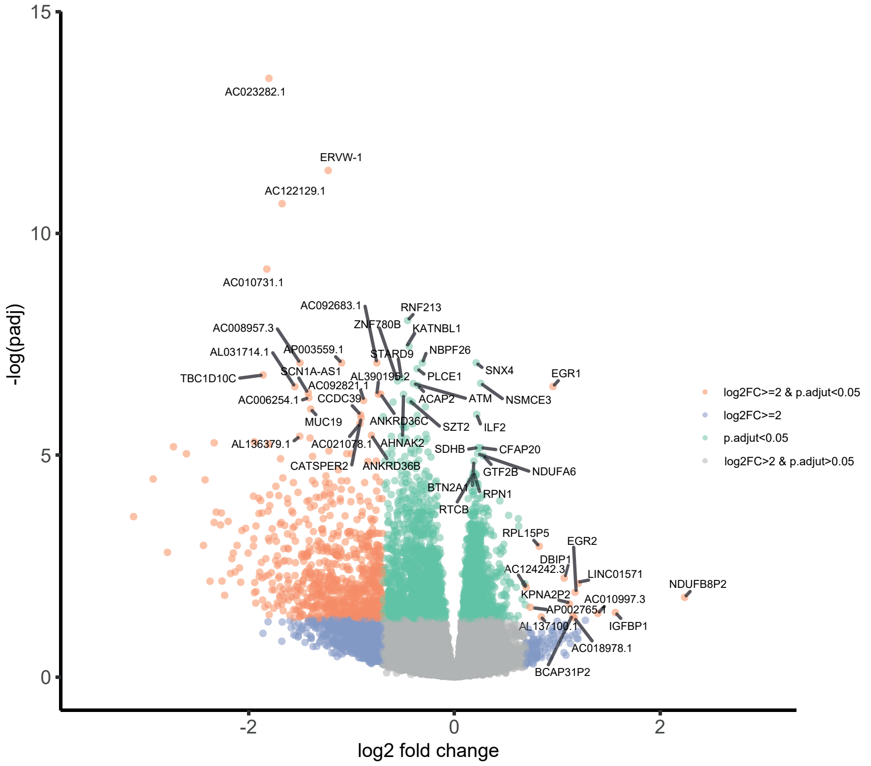


b

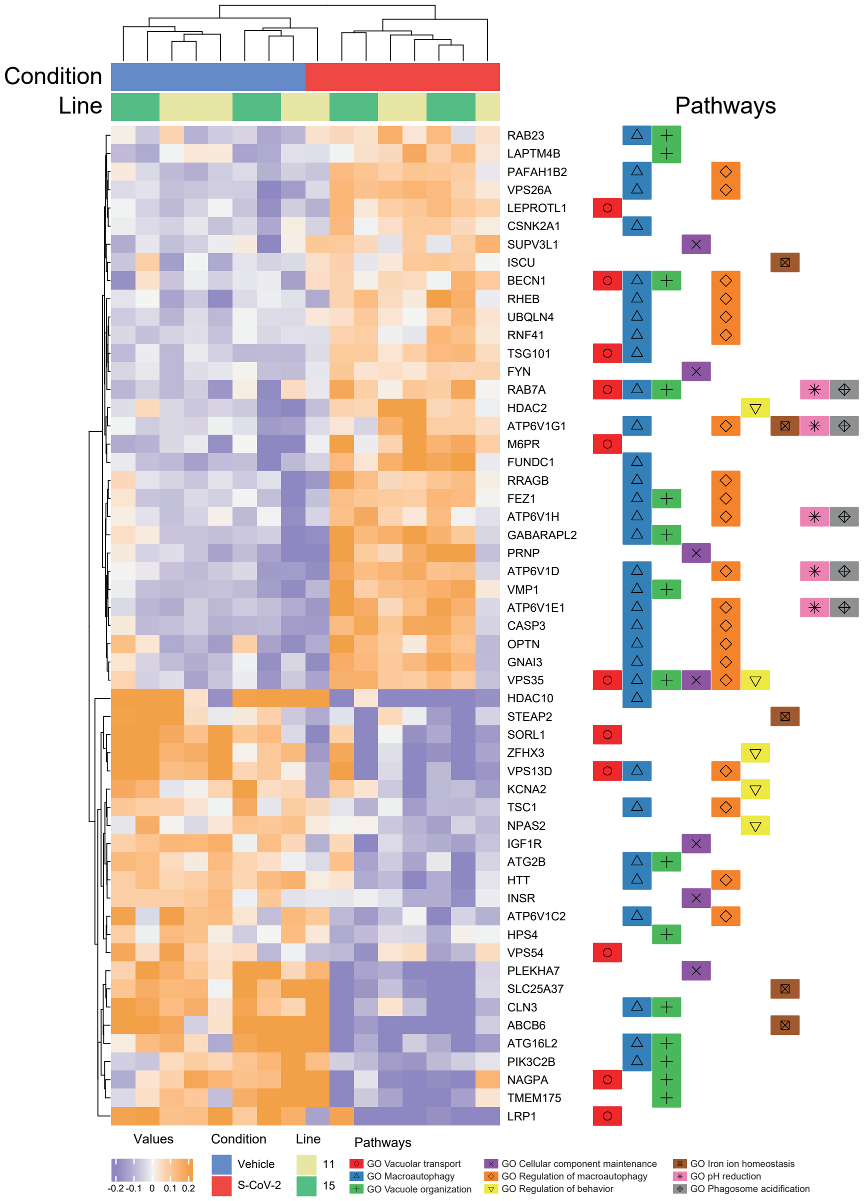

### Supplementary Figure 3

#### Extended data Figure 3

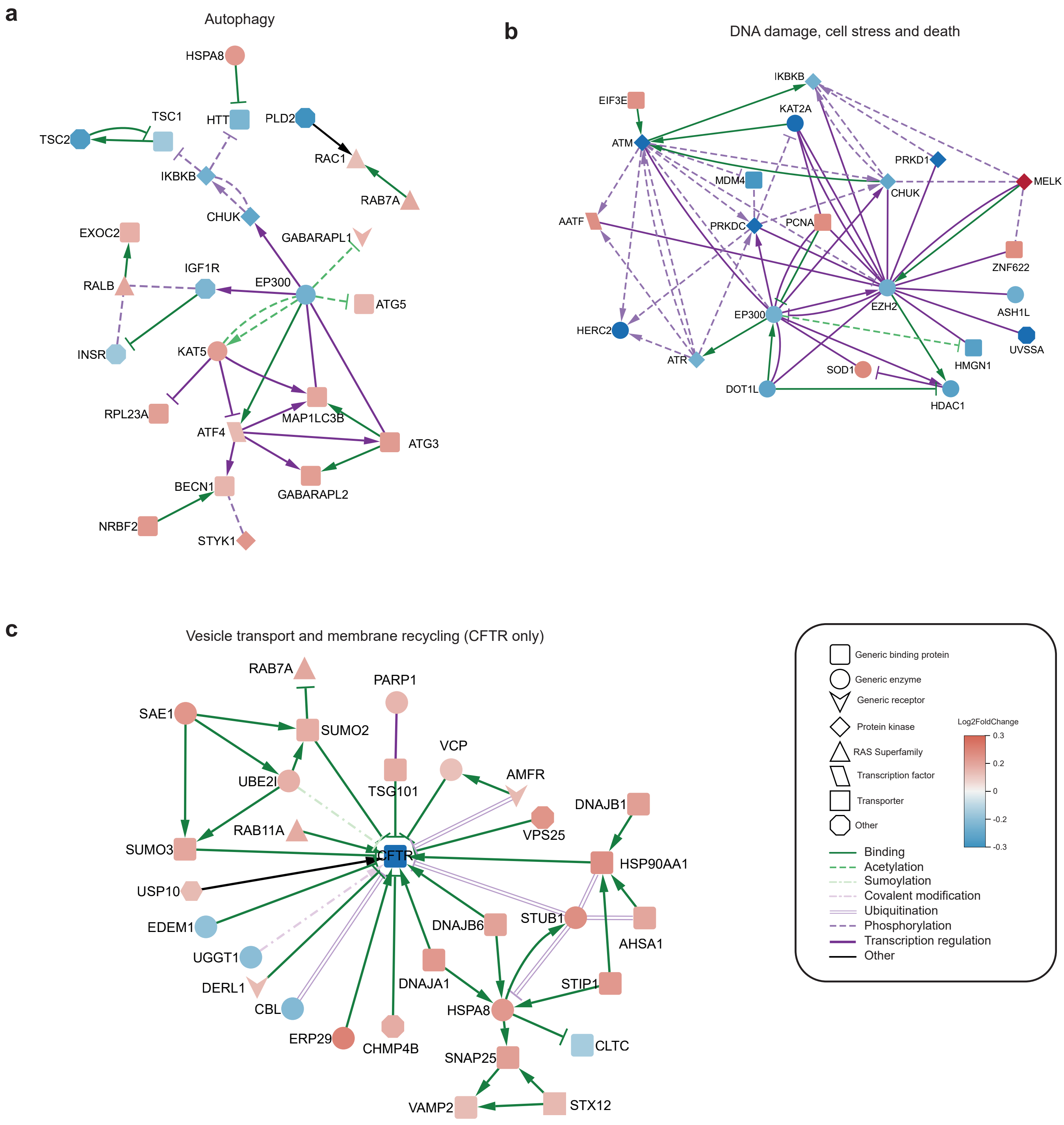

### Supplementary Figure 4

#### Extended data Figure 4

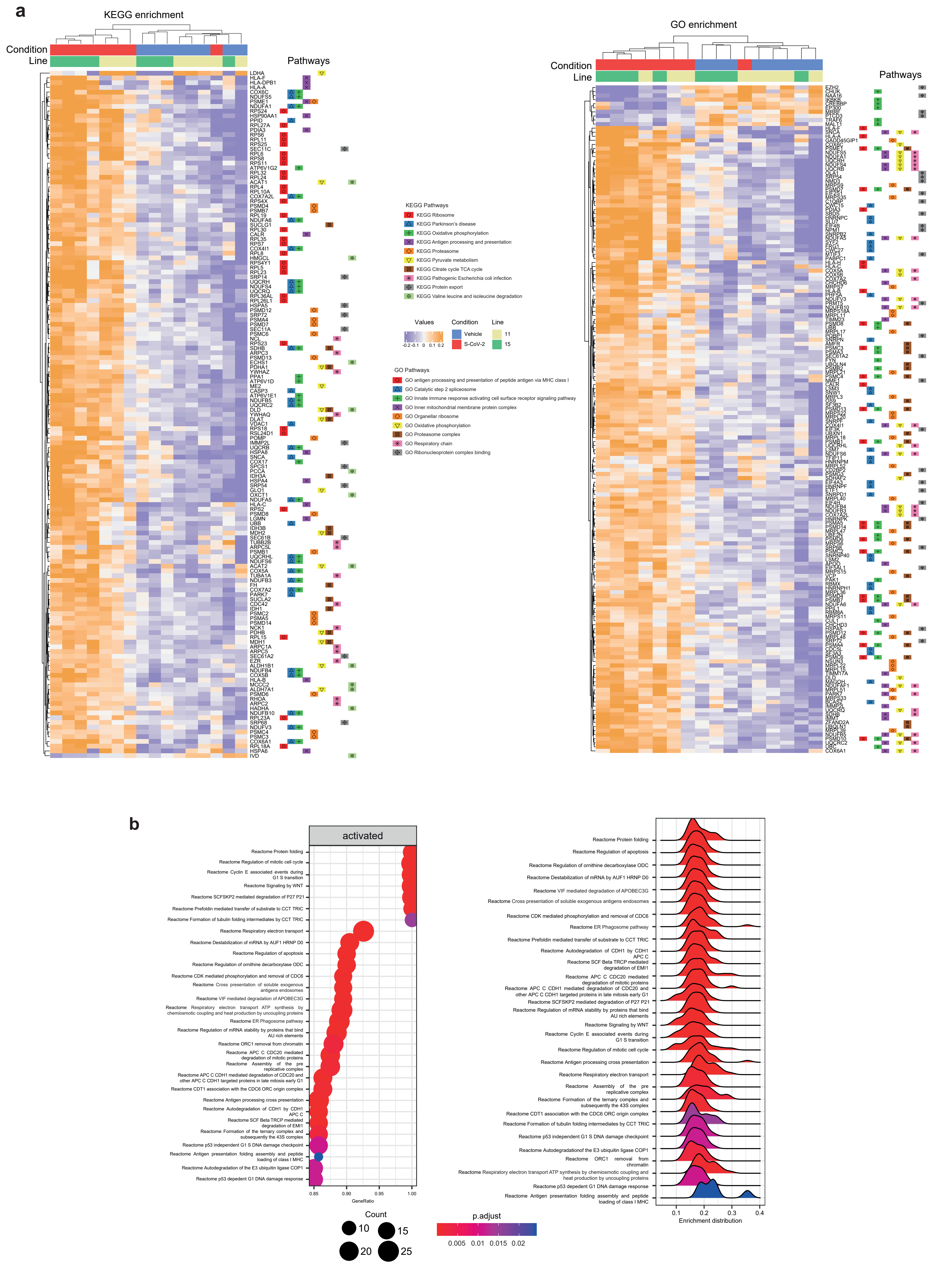

### Supplementary Figure 5

#### Extended data Figure 5

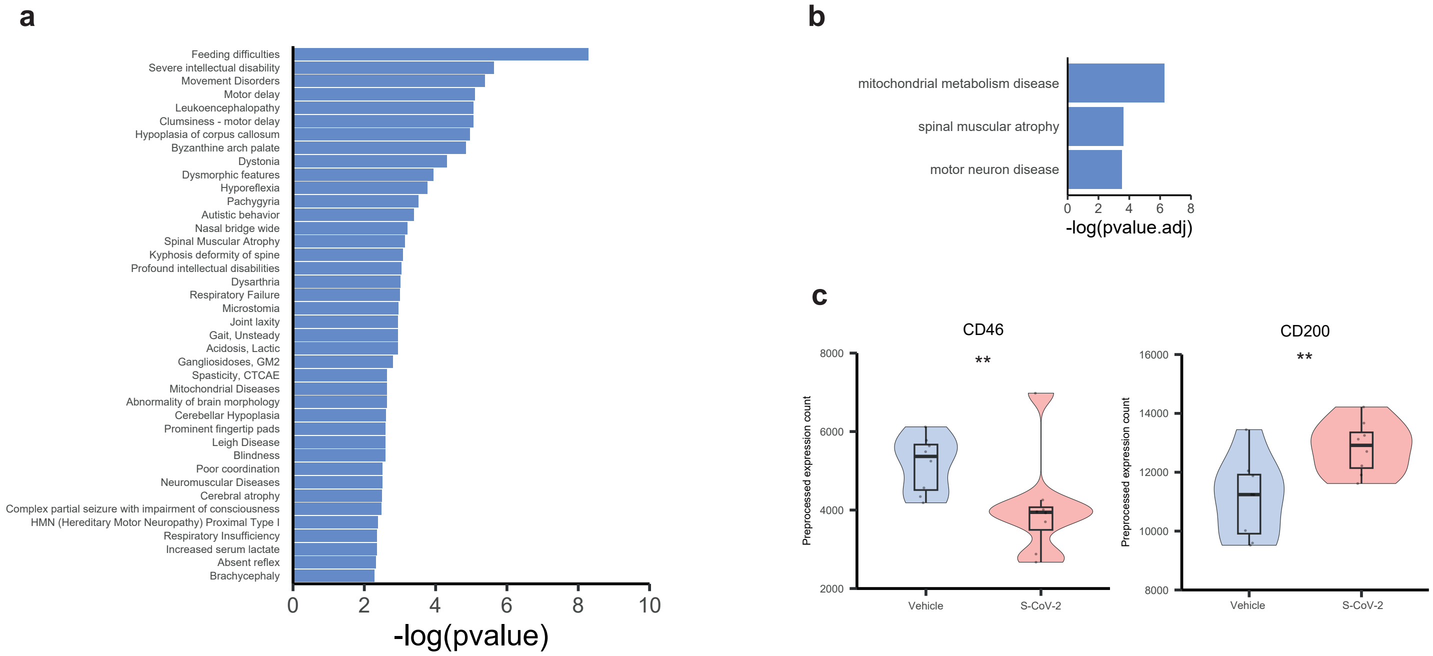

### Supplementary Figure 6

#### Extended data Figure 6

a

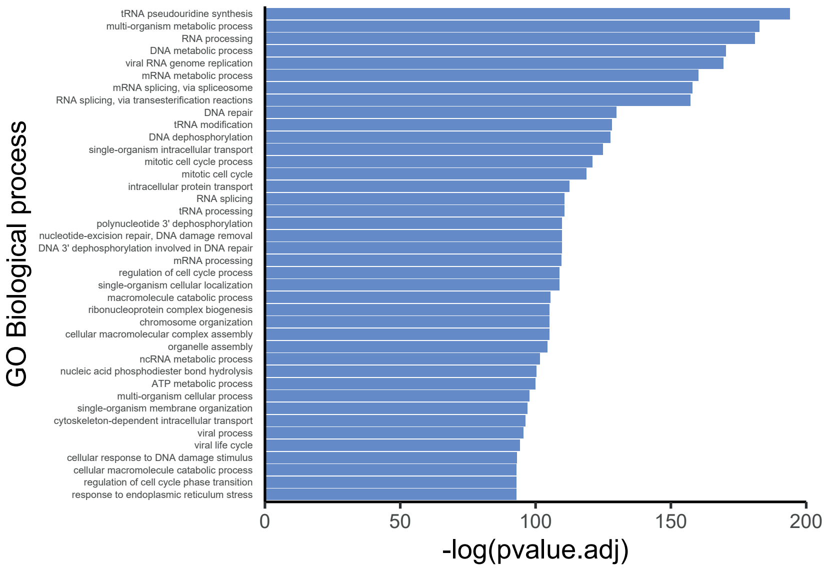


b

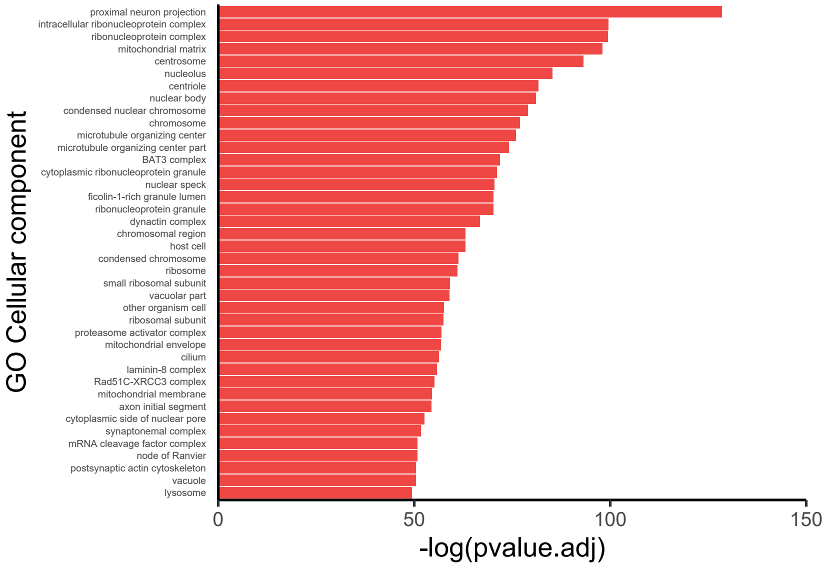


c

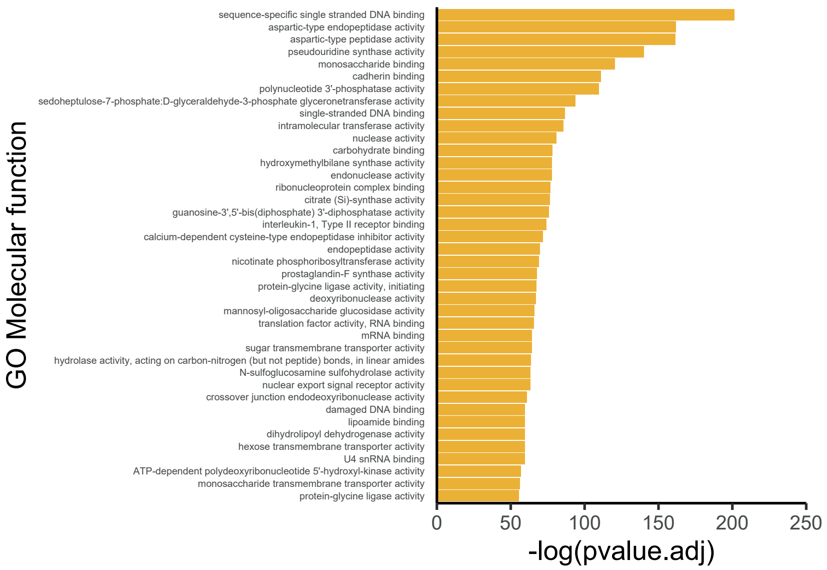
